# The Characteristic Timescales of the Arabidopsis thaliana Metabolism and the Phenotypic Effects of Forcing Them

**DOI:** 10.1101/2025.01.08.631846

**Authors:** Giovanni Santoro, Enrico Rolli, Ada Ricci, Ludovico Cademartiri

## Abstract

Response mechanisms preserve steady states in complex systems (including living systems). These mechanisms respond with characteristic timescales. Therefore, they can be identified and probed with oscillatory forcing. This approach is rarely used in plant biology.

Sub-circadian intermittent illumination (i.e., square waves of light/dark periods of equal duration) of *Arabidopsis thaliana* plants cause remarkable phenotypical effects. We show these effects are highly dependent on the duration of the light-dark cycle, identifying characteristic timescales in three distinct windows (10.5-42 s, 168-336 s, and 1350-5400 s). Among these effects is an extension of the life cycle by 60%, a 2-fold increase in flower setting, and a near complete elimination of lateral root development.

This oscillatory light forcing is compatible with vertical farming where it could reduce energy and chemical inputs and highlights the methodological value of temporal-forcing of environmental parameters on sub-circadian timescales in plant systems.

---

Any system that tends to return to an equilibrium or steady state (e.g., a tuning fork) takes some time to do so. Its inverse is the characteristic frequency of that response. When these response mechanisms (e.g., its elasticity) are tested by the appropriate stimulus (e.g., mechanical stimulation) that oscillates in intensity at a frequency close to its characteristic one (e.g., 440 Hz for a standard A tuning fork), it will respond very distinctly^1-3^. These strong responses take not only the form of resonances^4,5^, but also of inhibition^6^, synchronization^7^, ratcheting^8^, etc.…. But what do we learn from these responses?

Complex systems have a multitude of response mechanisms that keep them in a steady or, less commonly, an equilibrium state. Due to the feedback loops in which these mechanisms are often entangled, it can be hard to identify individual mechanisms. A specific oscillatory stimulus of a specific frequency can then amplify (and therefore isolate) a single mechanism out of all the ones present in the system. This is the fundamental premise of spectroscopy techniques in chemistry (e.g., infrared spectroscopy, nuclear magnetic resonance), which leverage the easily accessible high frequencies of electromagnetic radiation to probe the response times of atomic or molecular-scale systems (e.g., the C-H bond).

Larger scale systems, whether simple (e.g., a tuning fork) or complex (e.g., the biosphere), tend to respond at much longer timescales. Organisms (or organs within them) are essentially large-scale chemical networks that are self-regulated to maintain a certain steady state^9^. They are large, and extremely complex, so they have a multitude of response mechanisms to different stimuli^10^, each of which operate on different but probably not extremely fast timescales (largely because mass transport is often required). Based on this premise, our hypothesis was that temporal oscillations in light intensity could affect plant physiology (and therefore phenotype) by either amplifying or inhibiting metabolic pathways.

The use of oscillatory inputs to probe living organisms is much rarer and generally concentrated on systems that have a clear dynamic component (e.g., the heart^11-13^, the brain^14-16^). While a remarkable amount of work has been conducted to understand the impact that different wavelengths of light had on the metabolism of plants^17^, remarkably little exists in the literature about oscillations of light intensity, in spite of the well-understood importance of the circadian clock in regulating plant biology^18,19^. While some researchers have studied the characteristics of daylength perception and the developmental consequences (e.g. floral induction) following an alteration in the length of the light/dark period, correlating any changes with modifications in the expression of specific genes^18,20,21^, only few have studied the effect that different frequencies in the light/dark alternation can have on the development of the plant organism^22^. In this regard, in order to decrease the energy requirements for greenhouse lighting, Song et al. reported that Arabidopsis plants subjected to pulsed light with extended dark periods developed substantially similarly to control ones provided with 12 h light treatment^23^.

Our first *in vitro* experiments immediately showed a remarkable number and variety of strong (and, in some cases, unprecedented) phenotypic responses as a function of the specific cycle period of illumination (cf. Figure 1C).

**Figure 1.**
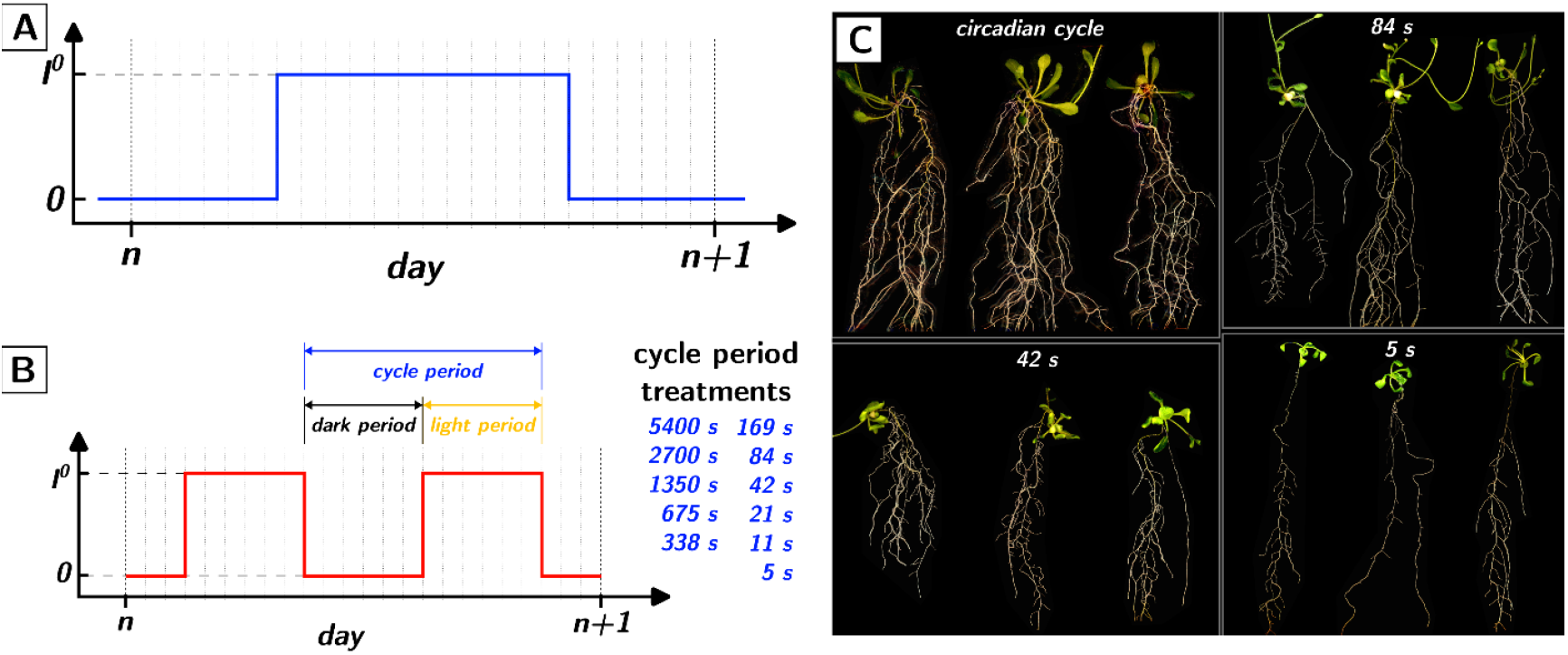
Subcircadian light intensity oscillations. **(A)** Circadian cycle of illumination described as a plot of light intensity as a function of time. **(B)** Subcircadian treatments used in this work, as square waves of illumination intensity with different total cycle periods. **(C)** Representative images of *A. thaliana* plants grown in Petri dishes under different cycle periods for 20 days, showing remarkable phenotypic adaptation.

We here summarize the main findings of our *in vitro* experiments (cf. Figure 2). As a summary we concentrate on the data at day 10, where most plants had not overgrown the plates and where phenotypes had sufficiently developed. All data for all days are provided in the Supporting Information. In all panels in Figure 2 the statistical significance between each pair of treatments is shown as colormaps (green for p<0.05). Peaks or troughs in the phenotype (i.e., phenotype of a treatment being significantly different that the two neighboring treatments) are identified by green crosses along the diagonal (outlined in blue in the figure). Discontinuities (i.e., phenotype of a treatment being significantly different that a single neighboring treatment) are instead identified by green “dumbbells” across the main diagonal (outlined in blue in the figure). As we detail below, all discontinuities, peaks, and troughs in the phenotypes occur in three windows of cycle periods: 10.5-42 s, 168-336 s, and 1350-5400s (marked as blue rectangles in the plots).

**Figure 2.**
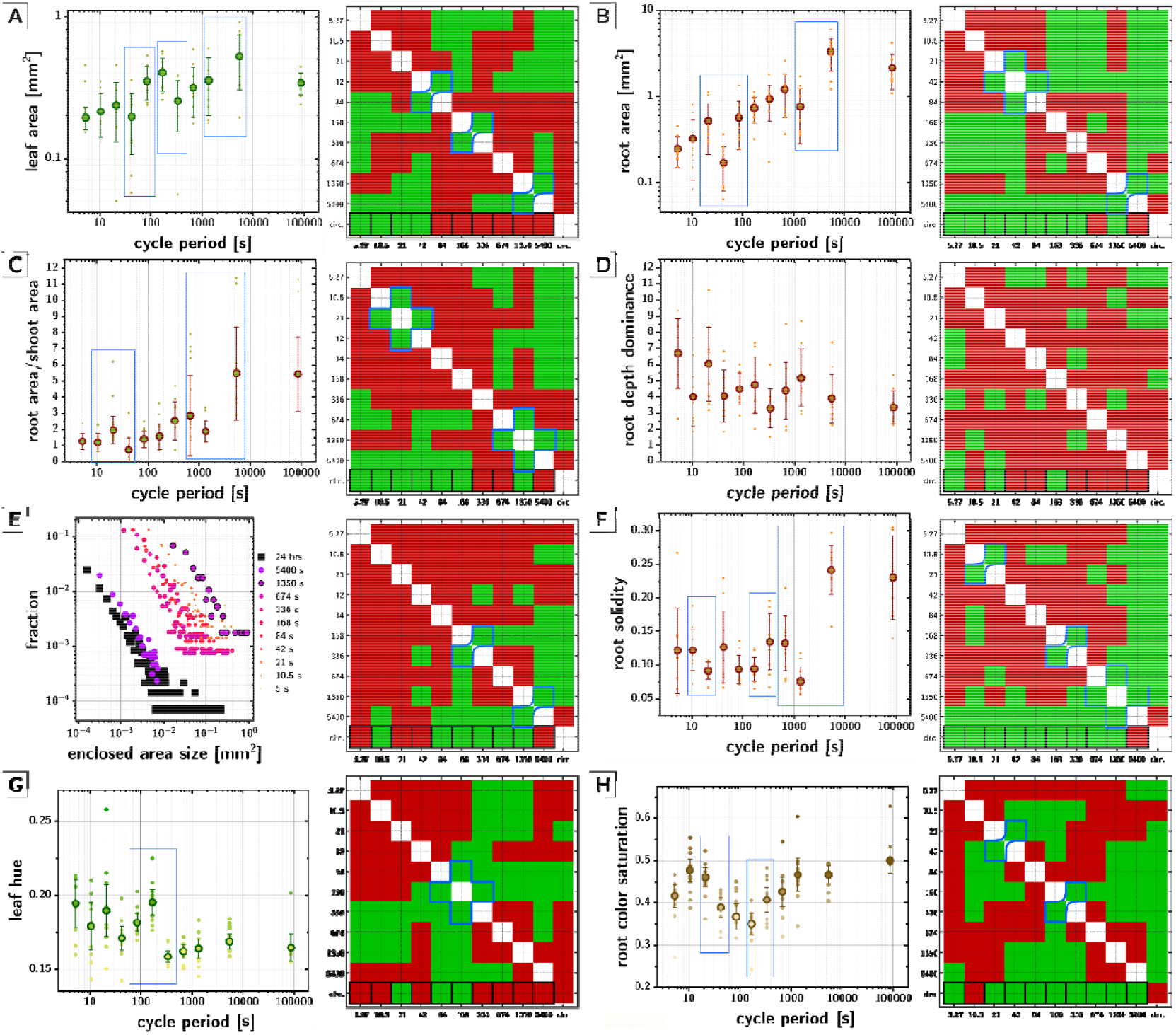
*In vitro* phenotyping of A. thaliana plants subject to sub-circadian light intensity oscillations (at day 10). In all plots, error bars are standard deviations. In each panel, colormaps indicate statistical significance (*green: p<0*.*05; t-test in case of datasets where normal distributions could not be excluded by an Anderson-Darling test or a Jarque-Bera test; Mann-Whitney U test in case normality could be excluded*) between each pair of treatments in the corresponding phenotype – outlined in black are highlighted the comparison with circadian control. In blue are highlighted the discontinuities in the phenotype’s dependency on cycle period. **(A)** Total leaf area as a function of the cycle period (loglog). **(B)** Total root area as a function of cycle period (loglog). **(C)** Root/shoot ratio (for areas) as a function of cycle period (semilog). **(D)** Root depth dominance (root depth/root width) as a function of cycle period (semilog). **(E)** Distribution of areas fully enclosed by roots as a function of cycle period (loglog) – data averaged over all growth days. **(F)** Root solidity (root area/root convex area) as a function of cycle period (semilog). **(G-H)** Leaf hue and root color saturation as a function of cycle period (semilog) – data averaged over all growth days.

The overall size of the plants (cf. Figure 2A-B) is significantly affected (at 10 days) by the cycle period. Compared to control, all cycles shorter than 84 s produced smaller total leaf areas, and all cycles shorter than 674 s produced smaller root areas. Root areas especially are reduced by one order of magnitude in the shortest cycle periods as compared to the circadian control. The trend of these phenotypes is not monotonic with the cycle period: a clear trough in root areas occurs at 42 s cycle periods which produced minuscule plants that seemed to struggle to survive; discontinuities occurred between 42-84 s, 168-336 s in the leaf area and 1350-5400s in both leaf and root area.

The root-to-shoot ratio (in area terms) also shows strong cycle period dependence (cf. Figure 2C). All cycles shorter than 336 s yielded significantly smaller root/shoot ratios than the control. A trough and a peak occur at 1350 s and 21 s respectively. Qualitatively, a large difference seems to occur between 1350 s and 5400 s (below 1350s root/shoot area ratio ∼2, while above 5400s is ∼5).

The architecture of the root system appears also to be influenced by cycle period as well (cf. Figure 2D). While these data at day 10 fail to convey the magnitude of the changes as growth proceeds, by day 20 the difference in lateral rooting for the shortest cycle periods is very significant (as shown in Figure 1C). The root depth dominance (ratio between depth and span of the root system) increases significantly as the cycle period is shortened: at 5 s cycle periods, roots are nearly devoid of lateral root, especially close to the shoot, producing a rather inverted vertical distribution of root biomass, as compared to circadian conditions.

The above observations on root architecture convinced us to verify whether the root enclosed areas distribution was affected by the cycle period. The distribution of areas enclosed by the root (the areas that are entirely surrounded by roots when grown on a flat surface) is a remarkably conserved root phenotype that appears to be basically independent of the overall size of the root system as well as a multitude of environmental conditions^24^. The distribution is expected to be describable by a power law, consistently with the fractal, self-similar characteristics of root systems^25^. As shown in Figure 2E, for the first time we observed a strong environmental adaptation of this phenotype, with, on average, drastically larger enclosed areas for subcircadian cycle periods. Interestingly the slope of the datasets in the loglog plots (i.e., the exponent of the power law) appears to be conserved.

To further confirm the significant alteration of the root architecture, we characterized its solidity (i.e., the ratio between root area and the root convex area) as a function of the cycle period (cf. Figure 2F). All cycle periods shorter than 5400 s give significantly less “solid” root systems than the circadian control. A trough occurs at 1350 s, and significant discontinuities occur at 10.5-21 s and 168-336 s. These root architecture modifications are further confirmed by the Euler and Haussdorff numbers (cf. Supporting Information).

Our first thoughts about these results were that the shortening of the cycle period could be described as a form of stress and that at least the most evident phenotypic adaptation (the decrease in plant size with decreasing cycle period) could be explained by that. Our first test of this hypothesis was to characterize the hue of the leaves to assess whether the treatments would induce an evident change in the leaf color, often associated with various forms of stress^26^. As shown in Figure 2G, the leaf hue actually seem to “improve” with decreasing cycle periods (as already quite evident in Figure 1C), with a clear discontinuity between 168 and 335 s cycle periods. Root color saturation (i.e., intensity of the root color) also seems to be impacted by cycle period (cf. Figure 2H). Most cycle periods below the circadian control yield significantly less colored roots than the control.

The consistent and exclusive occurrence of discontinuities in the *in vitro* phenotypes in three distinct windows of cycle periods is strongly suggesting of characteristic timescales of response mechanisms of the organism.

Since the *in vitro* experiments were not very consistent with a stress-based explanation for the observed phenotypes, and considering these observation on development, we hypothesized that at least some of the observed phenotypes could be the result of significant changes in the rate of development. This hypothesis was preliminarily supported by the observation of much stronger flower setting than the circadian control in the the 168 s treatment.

To verify this hypothesis we conducted *in vivo* experiments (cf. Figure 3) on two treatments corresponding to two of the windows of cycle periods corresponding to discontinuities in the *in vitro* phenotypes (5400 s and 168 s).

**Figure 3.**
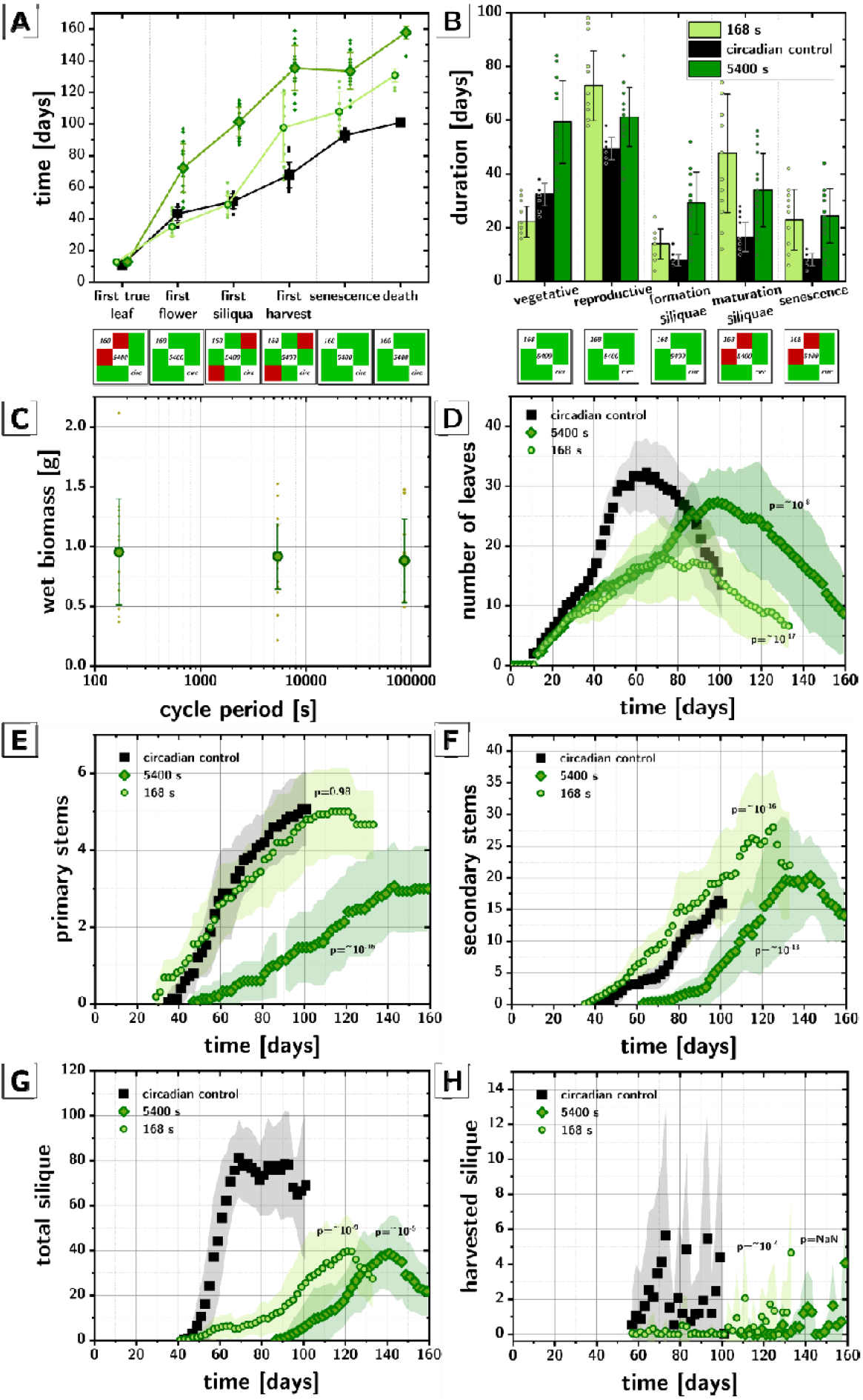
*In vivo* phenotyping of *A. thaliana* plants subject to sub-circadian light intensity oscillations. **(A)** Days since germination in which the life cycle phases initiate depending on the illumination cycle period. **(B)** Duration of the different life cycle phases depending on the illumination cycle period. **(C)** Fresh weight biomass collected from plants at the end of their life cycle as a function of the cycle period under which they were grown. **(D-H)** Number of leaves, primary and secondary floral stems, total siliquae produced, and harvested siliquae for *A. thaliana* plants grown under circadian (black squares), 5400 s (dark green rhombi), and 168 s (light green circles) cycle periods during their entire life cycle. Colored bands correspond to standard deviations (n=16).

The most obvious observation is that circadianly grown plants died after ∼100 days, while plants grown at 168 s and 5400 s lived for ∼130 days and ∼158 days respectively. The onset time of each developmental stage is drastically modified by subcircadian cycles (cf. Figure 3A) and, most importantly, each developmental phase is affected differently (cf. Figure 3B): the vegetative phase at 5400 s is ∼2 times longer than the control, while the one at 168 s is ∼30% shorter.

Consistently with our lack of consistent evidence of stress in the *in vitro* experiments, and in spite of the massive changes in phenotype, the fresh weight biomass at the end of the life cycle of each *in vivo* treatment was indistinguishable from control (cf. Figure 3C) again supporting the notion that sub-circadian illumination might not be necessarily a stressor.

The number of leaves over time (cf. Figure 3D) for the circadian control show a characteristic dome shape with a sharp increase during flower setting. In the 168 s treatment the dome shape is preserved but without the sharp increase after ∼40 days. The production of primary stems in the 168 s treatment is basically synchronous with the control, in spite of an earlier onset (cf. Figure 3E). This is somewhat the case also for secondary stems (cf. Figure 3F), where though the 168 s has a consistently larger production of stems and continues producing them for 30 more days, resulting in a total number about double that of the control. While flower production is amplified, the production and maturation of the siliquae in the 168 s treatment is stunted with only half as many siliquae produced (78 vs 39, cf. Figure 3G) and a third matured (18 vs 51,, cf. Figure 3H) when compared to control.

The 5400 s treatment shows instead a qualitatively analogous behaviour in leaf production (cf. Figure 3D) but drastically stretched in time, with sharp increase in leaf production not occurring until ∼70 days. Total number of leaves produced was not significantly different, even if the evolution in time was (p∼10^−8^). The production of primary stems in the 5400 s treatment was significantly stunted and delayed when compared to control (cf. Figure 3E), while for secondary stems, the production was similar to control but also delayed by more than a month. While the 5400 s treatment had fewer stems than the 168 s one, the number of siliquae produced was similar (cf. Figure 3G) but delayed by 20 days with respect to the 5400 s and by 70 days with respect to control. The maturation of the siliquae (cf. Figure 3H) was instead even more compromised than for the 168 s treatment with only 14 siliquae harvested per plant instead of 51 in the control.

Speculatively, one could suggest the extension of the lifespan could in part derive from a form of “starvation” of the plant, reminiscent of what happens in other organisms^27-30^. While on a 24hr basis, all treatments were exposed to the same number of photons, on a lifespan basis they were not, indicating a lower ability to use photons in the sub-circadian treatments. Potentially this could be due to the minimum timescale necessary to fully activate the production of photosynthate biochemically and/or phenotypically (e.g., leaf movement^19^).

## Notes

### Competing Interest Statement

The authors have declared no competing interest.

