## Supporting Information for "The Characteristic Timescales of the Arabidopsis thaliana Metabolism and the Phenotypic Effects of Forcing Them"

*Experimental design and methods*

**Plant choice.** We chose *A. thaliana* as a plant to test our hypothesis due to its status as model organism for small plants, short life cycle, and small size (therefore allowing us both in vitro and in vivo characterizations). The simplicity of the genome of *A. thaliana* would make it unlikely that, in case some effects would be found, these would be highly specific to the species.

**Culture.** We chose large plates (12 cm x 12 cm) and we planted a single plant per plate to avoid interference (either through plant-plant communication or in the determination of phenotypes by image analysis), and maximize the growth time (up to 20 days). Plates were grown upright under a LED lamp controlled through an Arduino controller to turn on and off according to a square wave with periods distributed logarithmically between 5 s and 43200 s (the circadian control) (cf. Figure 1A-B). In all cases, the lights were on for 50% of the time: the amount of photons impinging on plants in a 24 hr period was the same across all treatments. The plates were sealed with Micropore tape to limit CO_2_ depletion and accumulation^1^. The boxes were constructed of plywood (10 mm thick). The internal dimensions were 42x33x36 cm (WxHxD) and it was painted with anti-reflective black paint. A cooling fan (12 cm diameter) was placed on the back wall. The flow of air expelled by the fan was channeled into a double wall of the box, containing an optical labyrinth, so that the air could escape without letting light in. Likewise, the front opening was equipped with two flaps, one external and one internal, in order to prevent accidental entering light.

Germination was conducted in soil/agar (0.4%) with 0.25xMS medium to facilitate germination and transplant. On the third day the plants were transferred in the dishes with 1xMS 0.8% agar.

**Treatments.** Given the exploratory nature the work, we crafted the input illumination patterns using the following rationale: (i) all treatments should provide the same number of photons; (ii) all treatments should provide, as much as possible, all frequencies of electromagnetic (EM) radiation to ensure the observation of all EM-frequency-specific effects; (iii) treatments should cover the broadest possible range of frequencies of intermittence, compatibly with the capabilities of the equipment; (iv) the waveform had to be the simplest possible and compatible with the equipment; (v) the maximum of the waveform should correspond to optimal illumination, while the bottom of the waveform should correspond to darkness.

As a result, we used square waveforms (LED lights are not always compatible with dimming and their spectrum can change slightly with dimming), with an equal fraction of time on and off (cf. Figure 1A). We chose 10 periods (5400 s, 1350 s, 675 s, 338 s, 169 s, 84 s, 42 s, 21 s, 11 s, 5 s) in order to (i) span a wide range of cycle durations, (ii) contain an exact number of cycles within the 24 hr period (cf. Figure 1B). The 5s limit was dictated by the limitations of our LED lights. The circadian treatment (i.e., 12 hr of light and 12 of darkness) was used as control.

**Phenotypes.** We focused on those macroscopic phenotypes that could be collected with high accuracy, in a time-resolved manner, as much as possible *in vivo*. Therefore, our initial study in vitro (n=10), was conducted on plates that were scanned with a flatbed scanner once a day. Macroscopic phenotypes for the shoot and root system were then extracted from the images by image analysis.

**Image analysis.** The plates were scanned *in vivo*, so they could not be opened. Therefore, the scans were performed from the back. Consequently, the imaging photons from the scanner light had to travel through the gel twice. Due to the irregularities (e.g., bubbles, droplets, differences in thickness of the gel) and intrinsic coloration of the agar gel, it was necessary to subtract the background. We wrote a code the performs this task according to the following procedure (cf. Figure S1): (i) the raw image is first manually labeled to identify the outline of the roots and shoot; (ii) the code fills the unmarked spaces in the marked image (i.e., the background) with randomly positioned dots; (iii) the background color under each dot is extracted and averaged; (iv) the pointwise sampling of the background is then interpolated to create an image of the background; (v) the background is then subtracted from the raw image to obtain the normalized color of the plant; (vi) the manually marked areas were then used to segment the background-subtracted images into a shoot image and a root image.

The results of this protocol were very clean images that could be easily thresholded for phenotyping. In all cases, if the root of the plant was found to have touched the bottom of the plate, the plant was not considered in the phenotyping.

***In vivo* phenotyping.** As discussed later on, selected treatments from the *in vitro* experiments were repeated *in vivo* (n=15) to expose the plants to the light intermittence treatments for the entire duration of their life cycle. In this case, the macroscopic phenotypes that could be characterized *in vivo* (e.g., number of floral stems, number of leaves) were measured nearly every day, while the ones that would require sacrificing the plant (e.g., biomass) were only conducted at the end of the experiment.

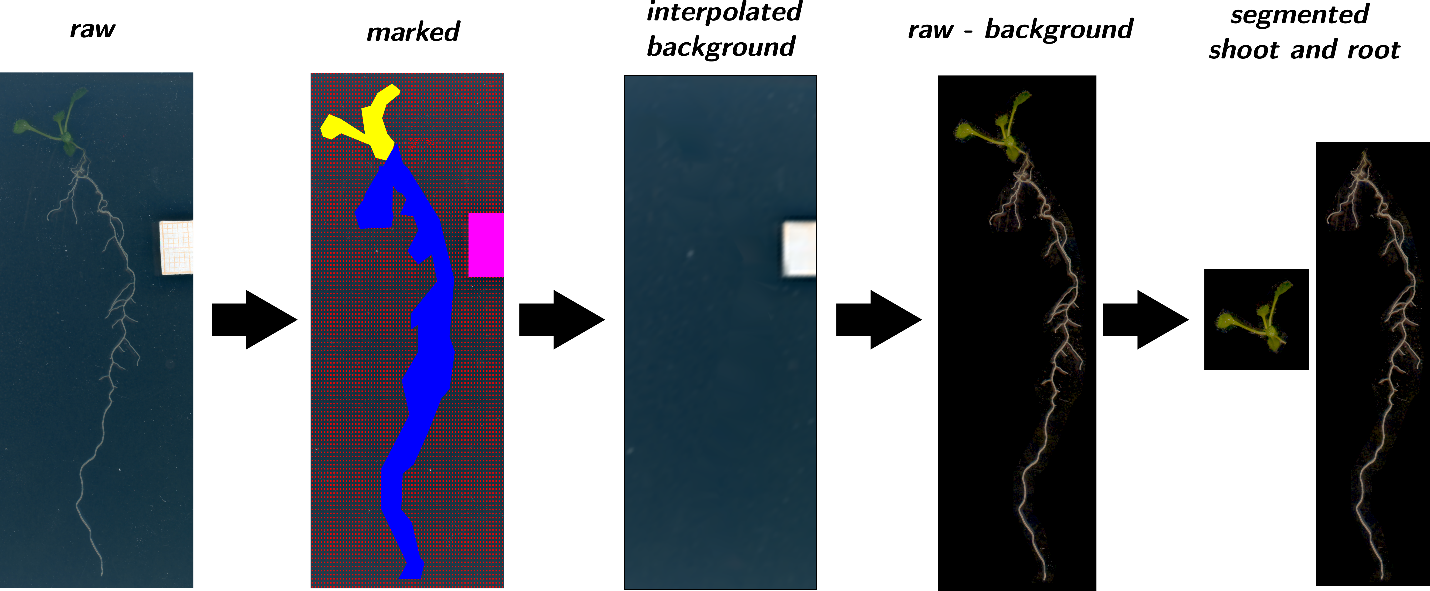

**Figure S1. Phenotyping protocol.** The raw scans were first marked to identify background area, root area and shoot area. The color values in the background area was interpolated from the sample points and subtracted from the original image. The image was then segmented to separate shoot and root.

*References*

1	Banerjee, S. *et al.* Stress response to CO2 deprivation by Arabidopsis thaliana in plant cultures. *PLoS One* **14**, e0212462, doi:10.1371/journal.pone.0212462 (2019).
